# From spiking neuronal networks to interpretable dynamics: a diffusion-approximation framework

**DOI:** 10.1101/2024.12.17.628339

**Authors:** Nosrat Mohammadi, Wilson Truccolo, Sydney S Cash, Anne-Lise Giraud, Timothée Proix

## Abstract

Modeling and interpreting the complex recurrent dynamics of neuronal spiking activity is essential to understanding how networks implement behavior and cognition. Nonlinear Hawkes process models can capture a large range of spiking dynamics, but remain difficult to interpret, due to their discontinuous and stochastic nature. To address this challenge, we introduce a novel framework based on a piecewise deterministic Markov process representation of the nonlinear Hawkes process (NH-PDMP) followed by a diffusion approximation. We analytically derive stability conditions and dynamical properties of the obtained diffusion processes for single-neuron and network models. We established the accuracy of the diffusion approximation framework by comparing it with exact continuous-time simulations of the original neuronal NH-PDMP models. Our framework offers an analytical and geometric account of the neuronal dynamics repertoire captured by nonlinear Hawkes process models, both for the canonical responses of single-neurons and neuronal-network dynamics, such as winner-take-all and traveling wave phenomena. Applied to human and nonhuman primate recordings of neuronal spiking activity during speech processing and motor tasks, respectively, our approach revealed that task features can be retrieved from the dynamical landscape of the fitted models. The combination of NH-PDMP representations and diffusion approximations thus provides a novel dynamical analysis framework to reveal single-neuron and neuronal-population dynamics directly from models fitted to spiking data.

## Introduction

A comprehensive understanding of the neural basis supporting behavior and cognition requires developing an interpretable appreciation of the intricate and complex dynamics that characterize neuronal network activity. This involves not only correlating neuronal activity with external stimuli and behavior but also modeling how past neuronal activity influences current spiking events. The use of nonlinear Hawkes processes with exogeneous inputs is an efficient statistical approach to model the recurrent and point process nature of neuronal spiking activity. Under some biologically relevant formulations, the models can be easily estimated via the (discrete-time) point-process generalized-linear-model framework [1–3]. Furthermore, these models can reproduce the large dynamical behavior variety of single spiking neurons, including canonical neuronal responses [4–6].

However, the direct dynamical analysis of estimated nonlinear Hawkes process models is a challenging issue due to their stochastic point process and non-Markovian nature. Theoretical studies have focused on determination of stability in variation for multidimensional nonlinear Hawkes processes [7–9], and mostly in specific cases that may be too restrictive for many neuroscience applications. Although other approximations have been proposed to analyze the stability of these models under exponential nonlinearities [10–13], the issue remains largely open. More importantly, beyond stability, the dynamics of these models, for both single-neuron and neuronal networks, have not yet been systematically investigated.

Here, we addressed this problem in two steps. First, we represented the single-neuron or neuronal network nonlinear Hawkes processes as an equivalent nonlinear-Hawkes piecewise deterministic Markov processes (NH-PDMPs) via the use of appropriate history kernels [2, 8, 9, 13], so that a set of stochastic differential equations with stochastic jumps (spikes) is obtained. Second, we applied a diffusion approximation to the jump processes. This diffusion approximation can work remarkably well even for single-neuron models [14]. Critically, these two steps allow for the use of analytical techniques from nonlinear dynamics, which offer quantitative and geometric insights into both the stability and dynamics of the nonlinear Hawkes processes. We demonstrated the accuracy of this approach by comparing it with exact continuous-time simulations of the corresponding NH-PDMP models, via specific algorithms we developed for this purpose. Our approach enables the identification of the dynamical repertoire of nonlinear Hawkes processes that reproduce the canonical responses of single-neuron activity, including tonic/phasic spiking and bursting, as well as type I and type II excitation. By applying this approach to neuronal networks, we show that complex winner-take-all and traveling wave dynamics are well captured by these NH-PDMP models. Finally, we apply our approach to human and nonhuman primate neuronal population recordings during speech processing and center-out reach movement tasks, respectively, and recovered within the dynamical landscape of the NH-PDMP models an organization of task features that are naturally present in the tasks.

## Results

We used nonlinear Hawkes processes with exogeneous inputs to model the dependence of a neuron’s instantaneous spiking rate on both the network’s past spiking history and external covariates, including stimuli and behavior [1]. At each time *t*, the conditional intensity function *λ* (*t*|ℋ _*t*_, *I*(*t*)) (called spiking rate thereafter) is conditioned on the neuronal spiking history ℋ_*t*_ up to time *t*, and exogeneous inputs *I*(*t*) (called inputs thereafter). Here, the spiking rate and the probability of observing a spike event for neuron *i* in a network of *J* neurons are given by (Figure 1(a))

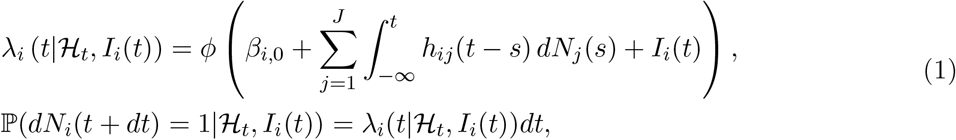

where *ϕ* is a nonlinear function and *β*_*i*,0_ is an offset parameter related to a background spiking rate. The term *N*_*j*_ (*t*) is the stochastic sample path counting the number of spike events for neuron *j* in the interval (−∞, *t*] with corresponding increments *dN*_*j*_ = 1 at each spike time and zero otherwise. The term *h*_*ij*_ is the time-causal history kernel capturing the spiking history effects. Specifically, *h*_*ii*_ is the auto-history kernel accounting for the past history effects on the same neuron *i*, while *h* _*ij*_ is the cross-history kernel accounting for the history effects from neuron *j* to *i* . We emphasize that the auto-history effects can go much beyond those present in renewal processes (where the neuron resets after a spike and only the time elapsed since the last event matters), as clearly seen in the adaptation dynamics in many neurons [15], for example. For notational convenience, henceforth we will write *λ*_*i*_ (*t*|ℋ_*t*_, *I*_*i*_(*t*)) as *λ*_*i*_(*t*).

**Figure 1.**
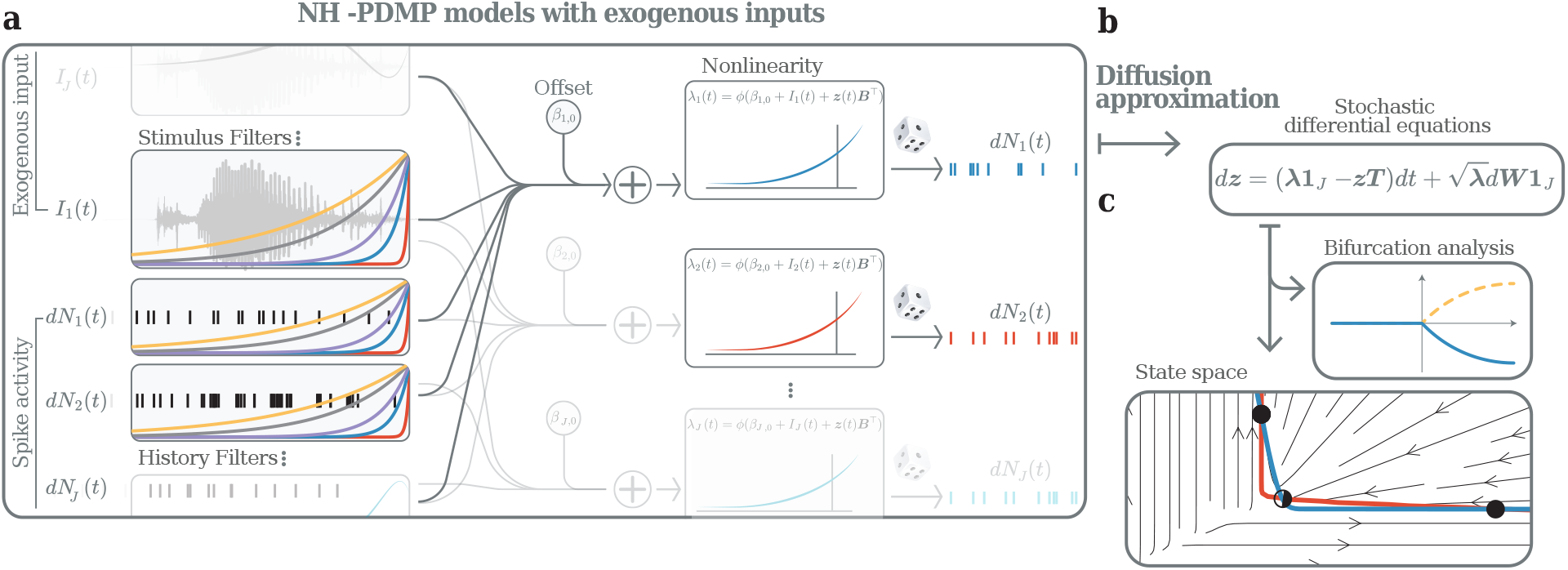
Nonlinear-Hawkes piecewise deterministic Markov process (NH-PDMP) models for spiking neuronal networks: (**a**) Schematic of a neuronal network based on NH-PDMP models with inputs. (**b**)The diffusion approximation transforms the SDE representation of the NH-PDMP into diffusion SDEs. (**c**)The dynamics of the diffusion SDEs can be analyzed qualitatively and quantitatively using dynamical system theory tools, such as state space and bifurcation analyses.

Determining the stability and dynamical system repertoire of the above single-neuron and neuronal network models based on nonlinear Hawkes process remains an important challenge. The challenge arises in part because of the discontinuous nature of the jump process *dN*_*i*_(*t*) reflecting the stochastic arrival of spikes. Most previous studies have focused on stability analysis, in particular stability in variation restricted to the case of Lipschitz nonlinearities [7, 9]. In neuroscience, on the other hand, the use of non-Lipschitz nonlinearities, specifically the exponential function, has several important experimental and statistical motivations (Methods). While some initial approximation results have been provided in this case, the issue remains open [10, 12, 13].

To address the above challenges, we approached the problem in two steps. First, we equivalently represent the nonlinear Hawkes process model by a NH-PMPD model. Second, we apply a diffusion approximation. These two steps allow transforming the nonlinear Hawkes process into a system of SDEs which can then be analyzed using standard dynamical systems tools. The proposed approach is general for any type on nonlinearity choice for the NH-PDMP models. Because its particular relevance to neuroscience as stated above, here we illustrate the approach within the case of exponential nonlinearities.

### Nonlinear-Hawkes PDMP (NH-PDMP) models

History kernels *h*_*ij*_ (*t*) can be approximated via special basis functions. In particular, when the history kernels are represented by sums of exponentials, or more generally Erlang functions [13, 16, 17], the nonlinear Hawkes process can be in principle equivalently represented as a NH-PDMP [2, 8, 9, 18]. NH-PDMPs are Markov processes that evolve deterministically in between stochastic jumps which occur at random time points, specifically neuronal spike times in our case. To benefit from the NH-PDMP formulation, we chose history kernels of the form 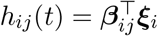, where ***β***_*ij*_ = (*β*_*ij*,1_, …, *β*_*ij*,*L*_)^*T*^ denotes the parameters for the *L* exponential functions 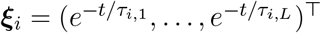 with corresponding time constants given by *τ*_*i*,*k*_. Additional time constants specific to cross-neuron interactions, e.g. *τ*_*ij*,*k*_ can be formulated too, but here, for simplicity, we focused on setting one time constant per neuron. Additional constraints are later formulated for data-driven models.

Based on the above history kernels *h*_*ij*_(*t*), eq. (1) is equivalently described by a NH-PMDP model given by the following system of stochastic differential equations (SDEs):

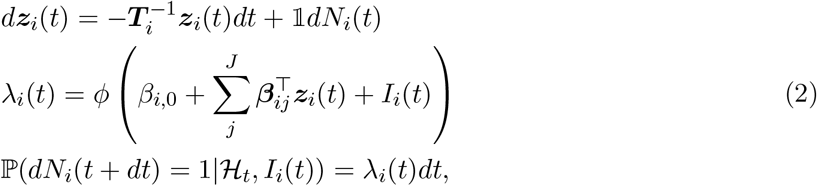

where ***z***_*i*_ = (*z*_*i*,1_, …, *z*_*i*,*L*_)^*T*^ is the activity supported by each history kernel of neuron *i* on its own time scale, ^𝟙^ = (1, …, 1)^*T*^ is a vector with *L* ones, and ***T***_*i*_ = diag(*τ*_*i*,1_, …, *τ*_*i*,*L*_). Spiking in the same and other neurons is influencing the dynamics of these variables through the weights ***β***_*ij*_ .

Exact continuous-time simulation of the process eq. (2) can be implemented by applying the time-rescaling theorem [1, 19, 20] while also taking advantage of the linear nature of the deterministic pieces of the PDMP representation (Methods). To our knowledge, the combination of time rescaling and the PDMP dynamics for exact continuous-time simulation, used throughout the manuscript, is new.

### Diffusion approximation

Next, we applied a diffusion approximation of the NH-PDMP process in both cases of single-neuron and network models. This is motivated by the known result that a jump process characterized by a Poisson process can be well approximated by a diffusion as the event rates grows larger and the jump sizes become correspondingly smaller [21, 22]. Previous studies have shown that this approximation can work remarkably well for single-neuron models [10, 14], especially if the spike events are not too sparse and the history kernels are smooth (Discussion). With the diffusion approximation, the jump process is effectively replaced by a Gaussian process which, by use of Itô’s lemma, can be written as [14]

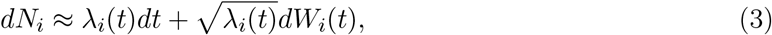

where *dW*_*i*_(*t*) corresponds to the increments of a standard Wiener process.

Using this approximation, we can now rewrite the NH-PDMP model (eq. (2)) as diffusion SDEs (Figure 1(b))

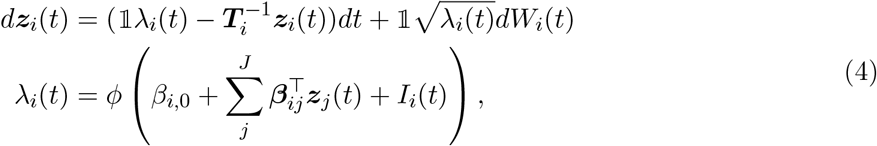

where the stochastic increments *dW*_*i*_(*t*) are independent across neurons. eq. (4) is a system of *J* · *L* SDEs. The critical advantage of this formulation is that the stochastic process is now a diffusion process, which facilitates its analysis via the use of moment closures [14, 23, 24] or other approaches. Because the diffusion SDEs only use continuous variables, akin to the frequently used rate models, the variable *λ*_*i*_(*t*) will thereafter be called the firing rate of the diffusion SDEs.

For convenience, we can rewrite the diffusion SDEs in a matrix form, where the *JL* state vector ***z***(*t*) = (*z*_1,1_, …, *z*_1,*L*_, *z*_2,1_, · · ·, *z*_*J*,*L*_)^*T*^ of the diffusion SDEs can be written as

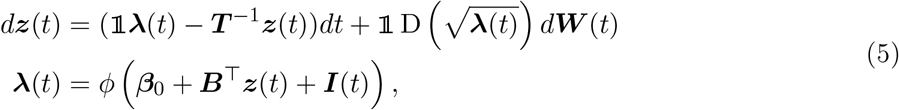

where the operator D(·) builds a diagonal matrix from a vector, ***T*** = diag(***T***_1_, …, ***T***_*J*_) is the block diagonal matrix of size *JL* × *JL* where each diagonal block is the matrix ***T***_*i*_, 𝟙 is the block diagonal matrix of size *JL* × *J* where each block is 𝟙 = (1, …, 1)^*T*^ with *L* ones, ***λ*** = (*λ*_1_, …, *λ*_*J*_)^*T*^, ***β***_0_ = (*β*_1,0_, …, *β*_*J*,0_)^*T*^, ***I***(*t*) = (*I*_1_(*t*), …, *I*_*J*_ (*t*))^*T*^, *d****W*** = (*W*_1_, …, *W*_*J*_)^*T*^, and ***B*** = ((*β*_11,1_, …, *β*_11,*L*_, *β*_12,1_, …, *β*_12,*L*_), …, (*β*_*J*1,1_, …, *β*_*JJ*,*L*_))^*T*^ is a matrix of size *JL* × *J*.

Although our approach is general with respect to the choice of nonlinearity in the NH-PDMP models, in this manuscript we will focus for experimental and statistical motivations (Methods) on the exponential nonlinearity *ϕ*(·) = exp(·). In the next sections, we analyzed the diffusion SDEs eq. (5) and showed how they accurately and geometrically predict the stability and dynamics of the original NH-PDMP models of single neuron and neuronal networks (Figure 1(c)).

### Stability and dynamics of single-neuron NH-PDMP models

We adopted two complementary approaches focusing respectively on the deterministic and stochastic stability of the diffusion SDEs approximated from the NH-PDMP models. In the first approach, we defined deterministic stability in the usual dynamical systems sense by examining the attracting and repelling properties of fixed points and related structures in the deterministic skeleton of the system. These fixed points (stable, unstable, saddle-node) can contribute to shaping the dynamics of the actual stochastic system, leading to metastability and changes in the properties of the stochastic flows in different regions of the state space, for example. In the second approach, we defined stochastic stability as the deterministic stability of first and second moment closures of the diffusion SDEs. In particular, bistability leads to metastability in the stochastic system.

We start by analyzing the stability properties of the diffusion SDEs of a single-neuron NH-PDMP model (eq. (4) with *J* = 1) using the first approach, i.e. by ignoring the stochastic term in eq. (4). The fixed points indicate the long-term dynamics of this system, and can be found analytically as a closed-form solution:

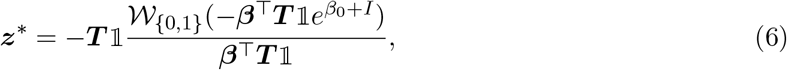

where 𝒲_{0,1}_(*x*) is the first or second branch of the Lambert function (see Methods, Figure 2(a)).

**Figure 2.**
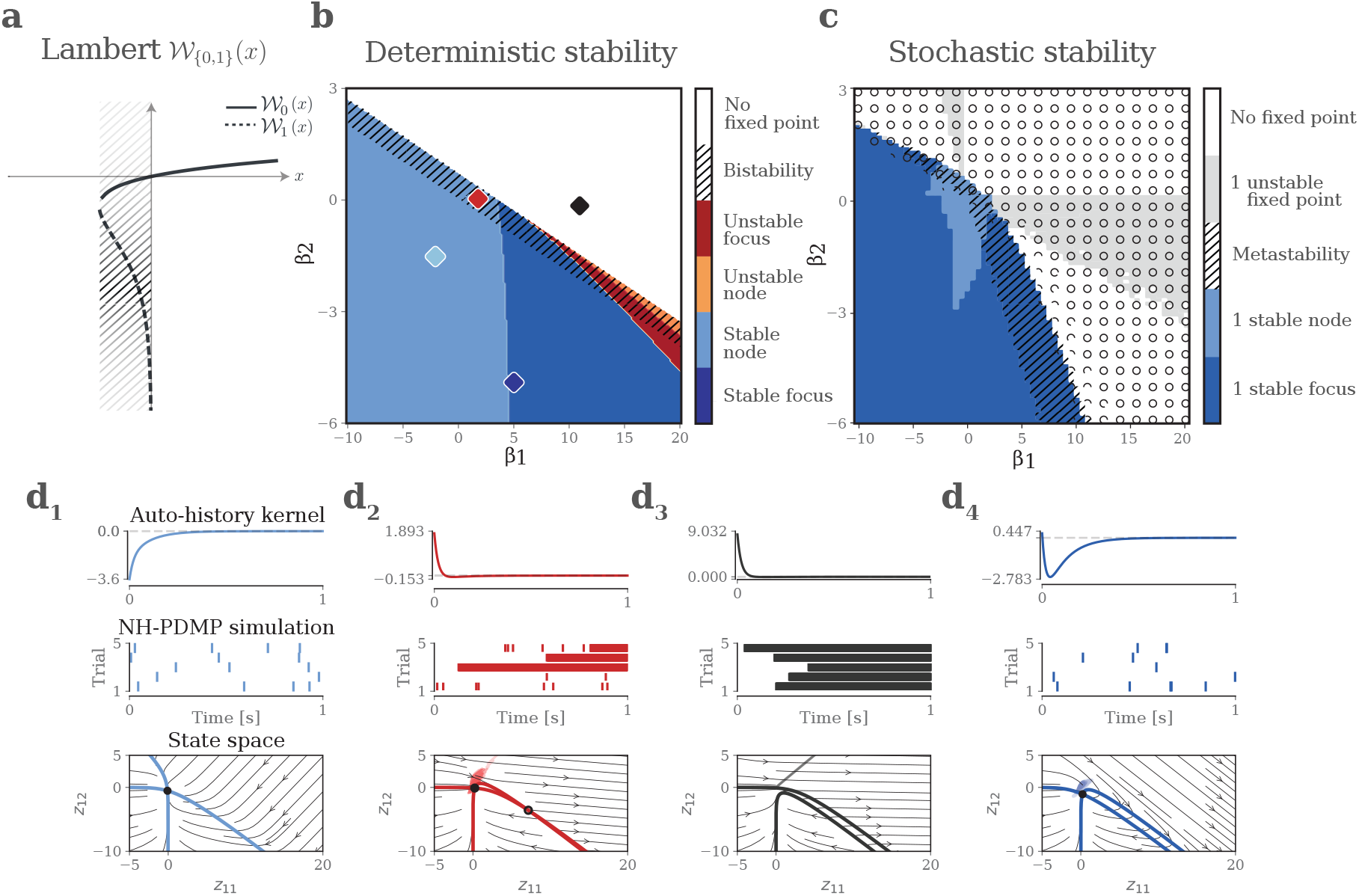
Stability of single-neuron NH-PDMP models: (**a**) Lambert W_*{*0,1*}*_(*x*) function. Hatched corresponds to the region with two real branches. (**b**) Parameter space of the deterministic part of the diffusion SDEs for a history kernel with two exponential functions with parameters *β*_1_ and *β*_2_, respectively. Colored regions indicate different dynamical regimes of existence and stability of the first fixed point obtained by analyzing the deterministic part of diffusion SDEs (eq. (4)). Hatch indicate the existence of an unstable second fixed point. Colored diamonds indicate different parameter choices for the examples shown in (d). (**c**) Parameter space of the moment equations of the corresponding full diffusion SDE. Colored regions indicate different stability regimes. Black circles indicate divergence of the conditional intensity (and number of spikes) in the exact continuous-time simulations of the NH-PDMP models (Eq. 2). The length of the circle indicates the number of initial conditions leading to diverging simulations (Methods). (**d**) Each column shows an example for different choice of the *β*_1_ and *β*_2_ parameters (colored diamonds in the parameter space: (1) *β*_1_ = −1, *β*_2_ = −1; (2) *β*_1_ = 2, *β*_2_ = −0.1; (3) *β*_1_ = −5, *β*_2_ = − 5; (4) *β*_1_ = 14, *β*_2_ = −1. First row: history kernels with two exponential functions defined by the above corresponding two parameters. Second row: 5 simulations of the corresponding NH-PDMP models (Eq. 2). Third row: state space spanned by the two variables *z*_1_ and *z*_2_ in the diffusion SDEs. Full circles indicate stable fixed points, colored lines are nullclines, and black arrowed lines show example trajectories.

The deterministic stability of these fixed points is determined by the Jacobian eigenvalues of the deterministic skeleton in the diffusion SDEs (see Methods). Although a closed-form solution cannot be obtained for these eigenvalues in the general case, several statements can be made from the analytical forms of the Lambert function and the Jacobian. This is summarized by the bifurcation diagram showing the existence and stability of long-term solutions (e.g. Figure 2(b)). If ***β***^*T*^***T*** 𝟙 *<* 0, i.e. below a hyperplane in the bifurcation diagram (colored regions below the hashed region in Figure 2(b)), the Lambert function is positive and only one fixed point exists, given by the branch _0_ (region with positive *x* in Figure 2(a)). Further, if all components of ***β*** are negative, this fixed point is stable (Figure 2(b), blue regions), otherwise stability can change depending on the parameters ***β, T***, and *I*. If *e*^*−*1^ *>* ***β***^*T*^***T*** 𝟙*e*^*I*^ *>* 0, i.e. above the previous hyperplane and below a second hyperplane (Figure 2(b), hashed region), two fixed points exist corresponding to the two branches 𝒲_0_ and 𝒲 _1_ of the Lambert function, whose stability can vary (hashed region in Figure 2(a)). Finally if ***β***^*T*^***T*** 𝟙*e*^*I*^ *> e*^*−*1^, i.e. above the second hyperplane, no fixed point exists and the system diverges (Figure 2(a), region with *x* smaller than the hashed values; Figure 2(b), white region).

To demonstrate the interest of the approach, we recovered the bifurcation diagram (Figure 2(b)) and its boundaries between different dynamical regimes for a history kernel with two exponential functions *β*_1_ exp(−*t/τ*_1_) + *β*_2_ exp(−*t/τ*_2_) (Methods). For negative values of *β*_1_ and *β*_2_, the system (Eq. 6) has one stable fixed point region (light blue region, example (1) in Figure 2(d) showing the state space and the geometry of the system’s dynamical properties. As described above, when ***β***^*T*^***T*** 𝟙 = *β*_1_*τ*_1_ + *β*_2_*τ*_2_ = 0, a bifurcation from infinity occurs, resulting in the creation of a new unstable fixed point, leading to the two fixed-point region of the Lambert function (hashed region, example (3) in Figure 2(d)). In this region of the parameter space, the system can diverge when its state goes beyond a certain range of values. Finally if (*β*_1_*τ*_1_ + *β*_2_*τ*_2_)*e*^*I*(*t*)^ *> e*^*−*1^, both fixed point disappear through a saddle-node bifurcation and the system diverges (white region, example (4) in Figure 2(d). For negative values of *β*_2_, if *β*_1_ increases, the stable fixed point of the upper branch of the Lambert function becomes a stable focus (eigenvalues of the Jacobian become complex, dark blue region, example (2) in Figure 2(d)). This stable focus leads to damped oscillations of the system with a specific period and damping factor. The stable focus then loses stability through a supercritical Hopf bifurcation to become an unstable focus (Figure 2(b), red region), and eventually an unstable fixed point (Figure 2(b), orange region). Increasing *β*_1_ even further leads to the collision of the stable and unstable fixed points through a saddle node-bifurcation, and a divergent system (Figure 2(b), white region).

So far, this analysis did not consider the stochastic nature of the diffusion SDEs. Stochasticity might affect the dynamics and stability of the models, as a large values of the noise term can lead to excursion away from a basin of stability. To analyze the full diffusion SDEs, we performed a moment expansion along with a Gaussian moment closure. The resulting equations are (Methods):

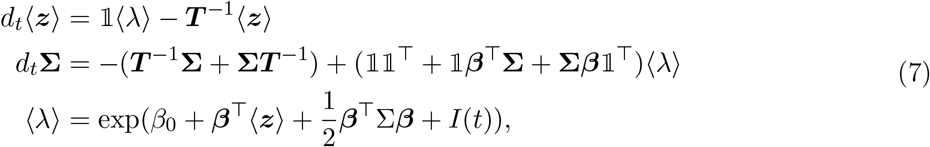

where ⟨***z***⟩ and **Σ**= ⟨ (***z***− ⟨***z***⟩)(***z***− ⟨***z***⟩)⟩^*T*^ are respectively the first moment and second central moment closures (covariance matrix) of the state vector ***z***. The fixed points of this system, as well as their linear stability as predicted by the Jacobian of the system, can then be evaluated numerically, leading to a modification of the borders between the different regimes (Figure 2(c)).

To assess the accuracy of the stability predictions based on both the deterministic and stochastic stability approaches applied to the diffusion SDEs, we compared them to exact continuous-time simulations of the single-neuron NH-PDMP models. For that purpose, we designed a specific algorithm based on a combined application of the time-rescaling theorem and the linear nature of the SDEs in the NH-PDMP representation (Methods). For history kernels consisting of two exponential functions, we found excellent agreement between the direct simulation of the NH-PDMP models (Figure 2(c), black circles) and the analysis of the moment-closure equations of the diffusion SDEs (Figure 2(c), colored regions). We reproduced the predicted stability ranges for different initial conditions. (The length of the black circles in Figure 2(c) corresponds to the number of different initial conditions that led to divergent simulations.) For some regions of the parameter space, simulations were always convergent (Figure 2(c), no circle), while for other regions they were always unstable (Figure 2(c), full circles). Finally, certain parameters choices led to divergent simulations only for some initial conditions (circular segment not forming a full circle). These regions correspond to regimes with two basins of attractions (hashed regions), where stochastic excursions can lead the system to cross the border of the basin of attraction.

In addition to these new stability analysis results in continuous time, our analytical results highlight the role of the initial conditions and the stochastic contributions in setting the different stability regimes, showing that for some regimes, different initial conditions or realizations may or may not lead to divergence in the systems. This dimension was ignored in the numerical simulations of the previous studies [10, 12, 13]. Previous studies also used discrete-time versions of nonlinear Hawkes processes, which we compared here as well as to our exact continuous-time simulation results. In these comparisons, we applied the diffusion approximation to the discrete-time nonlinear Hawkes process across the same parameter space for a history kernel with two exponential functions, following previous approaches [14]. As the time bin size decreases, the stability regimes of the parameter space increasingly better match the numerical simulations of the corresponding discrete-time nonlinear Hawkes processes. They however did not converge to the parameter space and exact continuous-time simulations of the NH-PDMP models, especially in the metastability regime.

### Dynamical repertoire of single-neuron NH-PDMP models

Nonlinear Hawkes process models have been shown to fit and reproduce a large variety of the dynamical repertoire of single neuron spiking activity [4, 5]. We asked whether the analysis method proposed here can reveal the dynamical repertoire of nonlinear Hawkes processes that support its dynamical flexibility. For this, we used Izhikevich single-neuron models under different parameter choices (table 1) to generate spike trains for canonical spiking behaviors, including tonic and phasic spiking, tonic and phasic bursting, and type I and type II neurons (Figure 3, Methods) [5, 6, 25]. For each case, we used multiple current step functions as input stimuli with variable amplitudes to generate a variety of responses for the training set. We then estimated NH-PDMP models using the (discrete-time) PPGLMs framework to fit the simulated spike train data. One or several exponential basis functions with different time constants *τ*_*k*_ were used for the history kernels (table 2). Next, we applied the diffusion approximation and determined the properties of the dynamics of the diffusion SDEs.

**Table 1:** Parameters for different spiking behaviors of the Izhikevich model.

| Spiking behavior | $a$ | $b$ | $c$ | $d$ | $I$ |
| --- | --- | --- | --- | --- | --- |
| Tonic spiking | 0.02 | 0.2 | -65 | 6 | 14 |
| Phasic spiking | 0.02 | 0.25 | -65 | 6 | 0.5 |
| Tonic bursting | 0.02 | 0.2 | -50 | 2 | 10 |
| Phasic bursting | 0.02 | 0.25 | -55 | 0.05 | 0.6 |
| Type I | 0.02 | -0.1 | -55 | 6 | 25 - 34 |
| Type II | 0.2 | 0.26 | -65 | 0 | 0.5 - 0.9 |

**Table 2:** Parameters of exponential kernels used for fitting different canonical response.

| Spiking behavior | $T[s]$ | $dt[s]$ | $\tau_1[s]$ | $\tau_2[s]$ | $\tau_3[s]$ | $\tau_4[s]$ |
| --- | --- | --- | --- | --- | --- | --- |
| Tonic spiking | 0.2 | 0.1 ms | 0.02 | NA | NA | NA |
| Phasic spiking | 3 | 0.1 ms | 0.02 | 0.1 | 0.5 | NA |
| Tonic bursting | 0.3 | 0.1 ms | 0.005 | 0.01 | 0.02 | 0.05 |
| Phasic bursting | 3 | 0.1 ms | 0.02 | 0.1 | 0.5 | 1 |
| Type I | 0.2 | 0.1 ms | 0.02 | NA | NA | NA |
| Type II | 3 | 0.1 ms | 0.02 | 0.1 | 0.5 | 1 |

**Figure 3.**
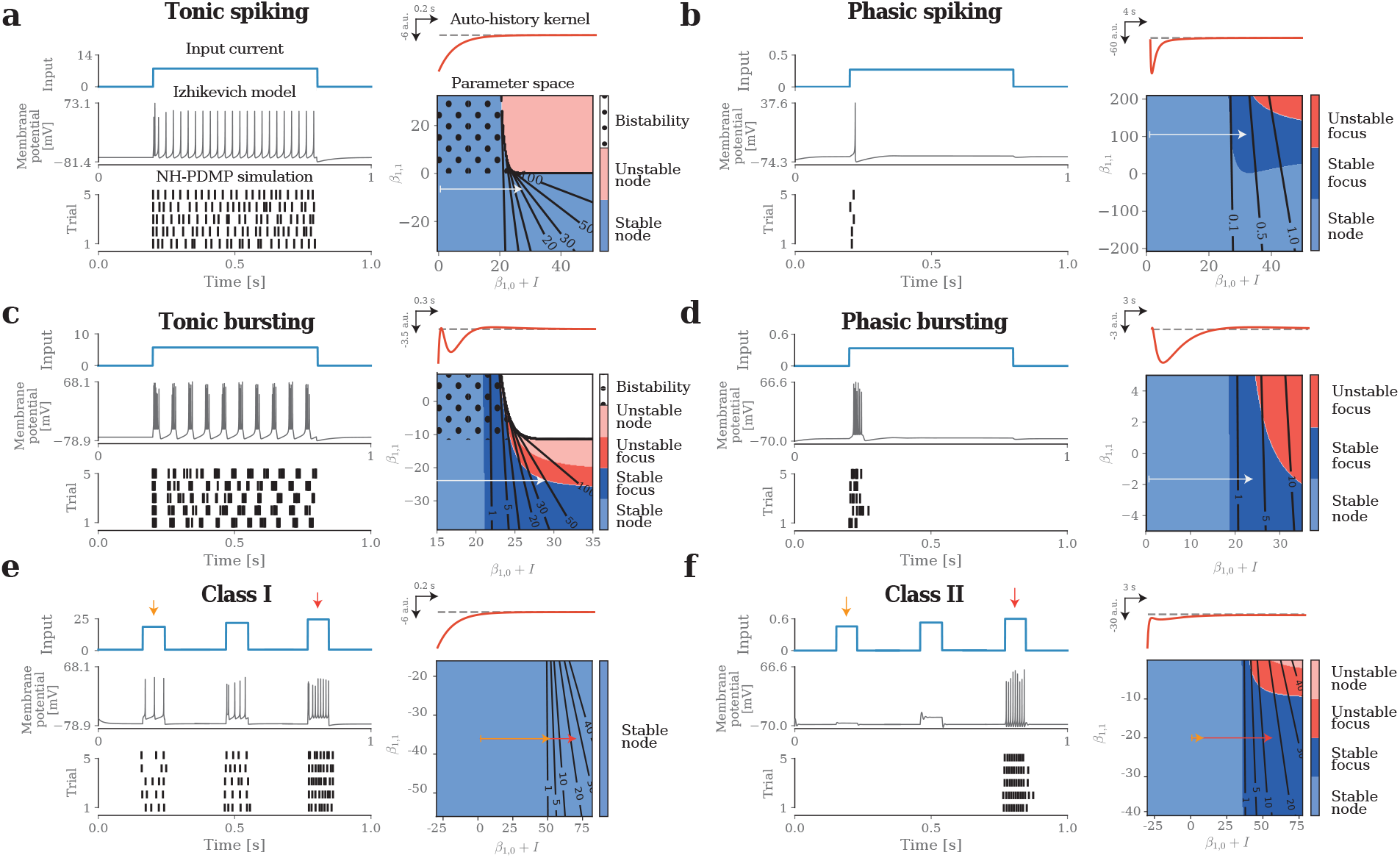
Dynamical repertoire of single-neuron NH-PDMP models: (**a**) Tonic spiking. Top left, example of a step function stimulus (blue). Middle left, corresponding Izhikevich neuron’s tonic spiking response (black). Bottom left, simulation of five trials of the fitted NH-PDMP model. Top right, history kernels of the estimated NH-PDMP model. Bottom right, parameter space of the deterministic part of the diffusion SDEs spanned by the input current and *β*_1_. Increasing the input *I*(*t*) for a fixed *β*_1_ leads to a transition along the horizontal arrow shown on the parameter space. Isoclines show the corresponding firing rate (in spikes/s) for the same neuron. The model transitions from a stable fixed point with very low rate to a stable fixed point with higher firing rate. (**b**) Phasic spiking. Same as in (a) for phasic spiking response. The model transitions from a stable fixed point to a stable focus regime with higher firing rate and high damping factor. (**c**) Tonic bursting. Same as in (a) for tonic bursting response. The model transitions from a stable fixed point to a stable focus with smaller damping factor and higher imaginary components, enabling the model to stay longer in the excited state. (**d**) Phasic bursting. Same as in (a) for phasic bursting response. The model transitions from a stable fixed point to a stable focus regime with higher firing rate and low damping factor. (**e**) Type I. Same as in (a) for type I response. The model transitions from a stable fixed point with very low firing rate to a stable fixed point with higher firing rate. The two horizontal arrows correspond to two input currents with different amplitudes. (**f**) Type II. Same as in A for type II response. The model transitions from a stable fixed point to a stable focus regime with higher firing rate and high damping factor.

We first examined tonic and phasic spiking regimes. On one hand, in the tonic spiking regime, a step input current results into sustained spiking activity in the Izhikevich model during the entire stimulation, and was captured using one exponential function for the history kernel (Figure 3(a)). The diffusion SDEs remains in the region of the parameter space with only one stable fixed point. The step input current simply moves the fixed point of the diffusion SDEs from a regime of low to high firing rate, thus resulting into the tonic spiking. This is confirmed by the examination of the parameter space of the moment equations of the full diffusion SDEs.

On the other hand, the phasic spiking regime of the Izhikevich model corresponds to a single spike emitted at the onset of the step current input, and was captured using three exponential functions (Figure 3(b)). In this case, the input current leads the NH-PDMP model to transition from a stable fixed-point regime with low firing rate to a stable focus regime with high firing rate and fast decay, allowing for only one spike to be emitted.

Next, we examined tonic and phasic bursting regimes. Tonic bursting in the Izhikevich model corresponds to sustained bursts of spiking activity, and was captured by using four exponential functions (Figure 3(c)). Similar to the phasic spiking regime, the diffusion SDE transitions from a stable fixed point with a very low firing rate to a stable focus, although with a small damping factor and higher rates. The small damping factor leads to many oscillations before the system finally sets into the fixed point, resulting in a quasi limit-cycle behavior and tonic bursting. For the phasic bursting regime, characterized by a single burst of spiking activity at the onset of the input current, we used relatively long time constants *τ*_*k*_ in the four exponential functions to capture the slow dynamics of the spike train data (Figure 3(d)). Similar to the tonic bursting behavior, the model transitions from a stable node with a very low firing rate to a stable focus with a higher firing rate. However, in contrast to the tonic bursting behavior, the model has now a damping factor strong enough to stop subsequent oscillations, preventing additional bursts to occur.

Finally, we investigated whether our analytical approach can uncover the dynamical repertoire of fitted NH-PDMP models that allow them to capture the type I and type II excitability in Izhikevich models. For this, we applied an input current with increasing amplitude to both neuron types. For type I neurons, characterized by a continuous increase of spiking activity for increasing inputs, we used a single exponential function in the history kernel. The diffusion SDE showed that the fixed point shifted from low to higher spiking rates for increasing currents, as expected for type I neurons (Figure 3(e)). Type II behavior, on the other hand, corresponds to an abrupt change of spiking activity for increased input current, and was captured by four exponential functions. Surprisingly, for models fitted to type II neurons, we did not find any switch from a fixed point of silent activity to a fixed point of high spiking rate (i.e., a bistable regime), as would have been expected. Instead, the model assigned a high value to the input current’s parameter, leading to a rapid (but not instantaneous) increase of the spiking rate from a stable node with very low rates to a stable focus with a high rate, leading to dynamics resembling the type II regime (Figure 3(f)). We also note that the fitted history kernels were similar to the ones obtained in previous studies [5]. We conclude that although the fitted NH-PDMP models can successfully reproduce the spiking response of Izhikevich neuronal models, they do not clearly differentiate between type I and type II neurons.

These results based on the diffusion approximation highlight an important aspect of the properties of the NH-PDMP models. The variables ***z***(*t*) in the diffusion SDE describe a temporally diffuse (rate) approximation of the NH-PDMP dynamics. Unlike the dynamics and bifurcations of the membrane potential activity of the modeled Izhikevich neurons, the corresponding parameter space reveals the firing rate dynamical landscape of the neuron. For instance, while the transition into tonic spiking regime corresponds to a saddle-node bifurcation in the Izhikevich model, it is captured by a simple translation of the same fixed point in the diffusion SDE. Overall, as revealed by our analytical approach, the diverse dynamics of diffusion SDEs in different regions of the parameter space directly reveals the spiking dynamical repertoire of the modeled neuron, rather than its underlying membrane potential dynamics.

### Stability and dynamics of neuronal-network NH-PDMP models

Our approach can be extended to explain the dynamical regimes of nonlinear Hawkes processes modeling the spiking activity of neuronal networks (Figure 4). For *J >* 1 neurons, no closed-form of the system’s fixed points exists for the deterministic part of the diffusion SDEs (eq. (5)). Fixed points can nevertheless be found numerically by solving ***ż*** = 0, and their stability and local dynamics can be determined by computing the eigenvalues and eigenvectors of the corresponding Jacobians. This analysis can be extended to the moment equations of the full diffusion SDEs (Methods):

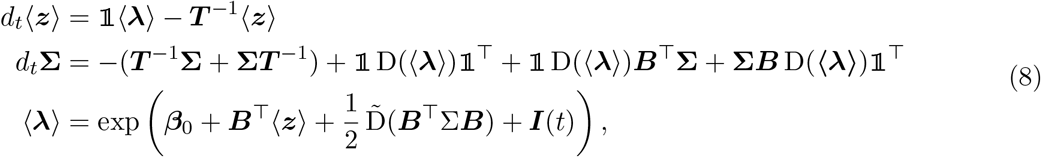

where the operator 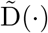 extracts the diagonal of a matrix as a vector. The fixed points of this system, as well as the linear stability predicted by the corresponding Jacobians, can then be evaluated numerically.

**Figure 4.**
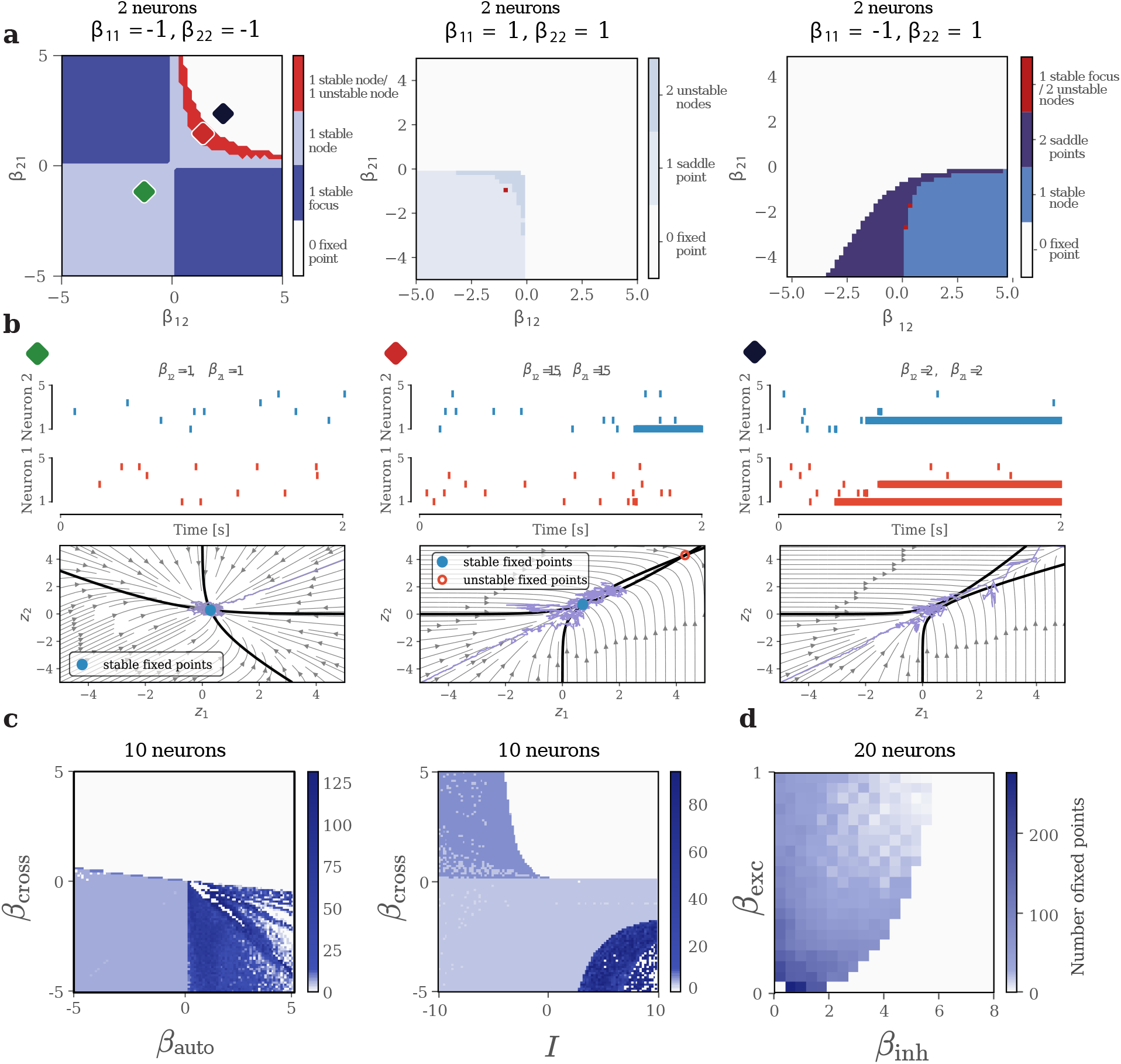
Stability of neuronal-network NH-PDMP models: (**a**) Parameter spaces of the deterministic part of the diffusion SDE for a network of two NH-PDMP model neurons. Each auto- and cross-history kernel was defined by one exponential functions with parameters *β*_*ii*_ (top) and *β*_*ij*_ (axis) respectively. Colored regions indicate different stability regimes. Colored diamonds indicate different parameter choices for the examples shown in B. (**b**) Top: spike trains from 5 different simulations of the corresponding NH-PDMP models. Bottom: State spaces spanned by the two variables *z*_1_ and *z*_2_ in diffusion SDEs for the different parameter values indicated by the diamond in the parameter spaces in A. Blue lines: trajectories in the state space; thick black lines: Nullclines; full circles: stable fixed points; arrows: system’s gradient. Parameter values indicated by the corresponding diamonds shown in the left parameter space in A. (**c**) Left: parameter space of the deterministic part of the diffusion SDEs for a network of ten neurons with a homogeneous connectivity (Methods) and varying parameters for the auto- and cross-history kernels. Right: same plot for varying inputs and parameters for the cross-history kernel. (**d**) Parameter space of the deterministic part of the diffusion SDEs for a network of 20 neurons with 80% excitatory neurons and 20% inhibitory neurons for varying inhibitory and excitatory parameters for the cross-history kernels.

As a demonstration of the approach, we analyzed the diffusion SDEs for two neuronal-network NH-PDMP models. First, we considered *J* = 2 coupled neurons and chose the history kernels to consist of a single exponential function. Such models already show a large variety of dynamical regimes. To better characterize these regimes, we systematically varied the cross-coupling parameters *β*_*ii*_ for different values of auto-history parameters *β*_*ij*_. For *β*_11_ = −1 and *β*_22_ = −1, both neurons have inhibitory auto-history. The system is bistable or unstable only when both coupling parameters are positive (Fig. Figure 4(a) left). For *β*_11_ = 1 and *β*_22_ = 1, both neurons have excitatory auto-history, restricting the part of the parameter space with saddle-point dynamics (i.e. with stable dynamics in some directions of the state space, and unstable dynamics in other directions) to the region defined by negative cross-coupling parameters (Fig. Figure 4(a) middle). Finally for *β*_11_ = −1 and *β*_22_ = 1, the second neuron is self-exciting, and the system is only stable when the second neuron inhibits the first neuron, with a large region of the parameter space corresponding to a bistability regime (Fig. Figure 4(a) right).

Second, larger network analysis showed that regimes with many fixed points can develop in some regions of the parameter space. We modeled a network of *J* = 10 neurons (with one exponential function for each auto- and cross-history kernels) such that all auto- and cross-coupling parameters are equal global auto- and cross-coupling parameters, respectively, i.e. *β*_*ii*_ = *β*_self_, ∀*i* ∈ [1, · · ·, *J*], and *β*_*ij*_ = *β*_cross_, ∀*i, j* ∈ [1,, *J*]^2^, *i* ≠ *j*. We then systematically increased the global auto- and cross-coupling parameters independently (Figure 4(d) left), as well as the input *I*(*t*) projecting equally to all neurons (Figure 4(d) right). We observed rich dynamics for inhibitory auto-coupling and excitatory cross-coupling, with a large number of fixed points.

Finally, we applied our analysis to a more biophysical network model including excitatory and inhibitory couplings for the cross-history kernels. We chose a network of *J* = 20 neurons with 80% excitatory and 20% inhibitory uniform coupling parameters with random connectivity pattern (Figure 4(d) right), modulated by global coupling parameters *β*_exc_ and *β*_inh_, respectively. We systematically varied these global coupling parameters. We observed that a high level of global excitation or inhibition will reduced the number of possible fixed points, suggesting that an appropriate balance of excitation and inhibition must be chosen to maintain rich dynamics.

### Dynamical repertoire of neuronal-network NH-PDMP models

We next asked whether our analysis approach can also help explain realistic biophysical circuit dynamics. As examples, we chose winner-take-all and propagating wave dynamics in network of neurons. These two types of dynamics are often explored in computational simulations of neural networks as well as actual population recordings [26, 27]. We simulated networks of Izhikevich neurons displaying winner-take-all and propagating wave dynamics using different input stimuli. As above, we then estimated NH-PDMP models using the PPGLMs framework to fit the spike trains of the simulated neuronal networks. Once the models were fitted, the diffusion approximation was applied and the corresponding diffusion SDE were analyzed.

For the winner-take-all network, we simulated two Izhikevich neurons with inhibitory cross-coupling functions in a regular spiking regime (Figure 5(a)). Each neuron received one of two distinct inputs taking different values (Figure 5(b) middle). A large difference between the two inputs led the neuron with the largest input to fire, while the activity in the other neuron was largely attenuated (Figure 5(b) top). Small differences between the two inputs could trigger either neuron to fire, depending on the stochastic realizations. The NH-PDMP models, estimated using three exponential functions, similarly displayed these two different types of dynamics (Figure 5(b) bottom). These two types of dynamics were captured by the diffusion SDE approximating the estimated NH-PDMP models. For inputs with large differences, only one fixed point exists in the state space (Figure 5(c), red and blue diamonds). Additionally, the nullclines are almost square angled, ensuring that when one input increases (i.e., when one nullcline moves with respect to the other nullcline), the resulting fixed point corresponds to a high firing rate for one neuron only (Figure 5(d), left and middle of the state space). By contrast, for small differences between inputs, three fixed points exists, corresponding to a bistability regime for which the winning neuron is determined by stochastic fluctuations (Figure 5(c) green diamond, and Figure 5(d), right state space).

**Figure 5.**
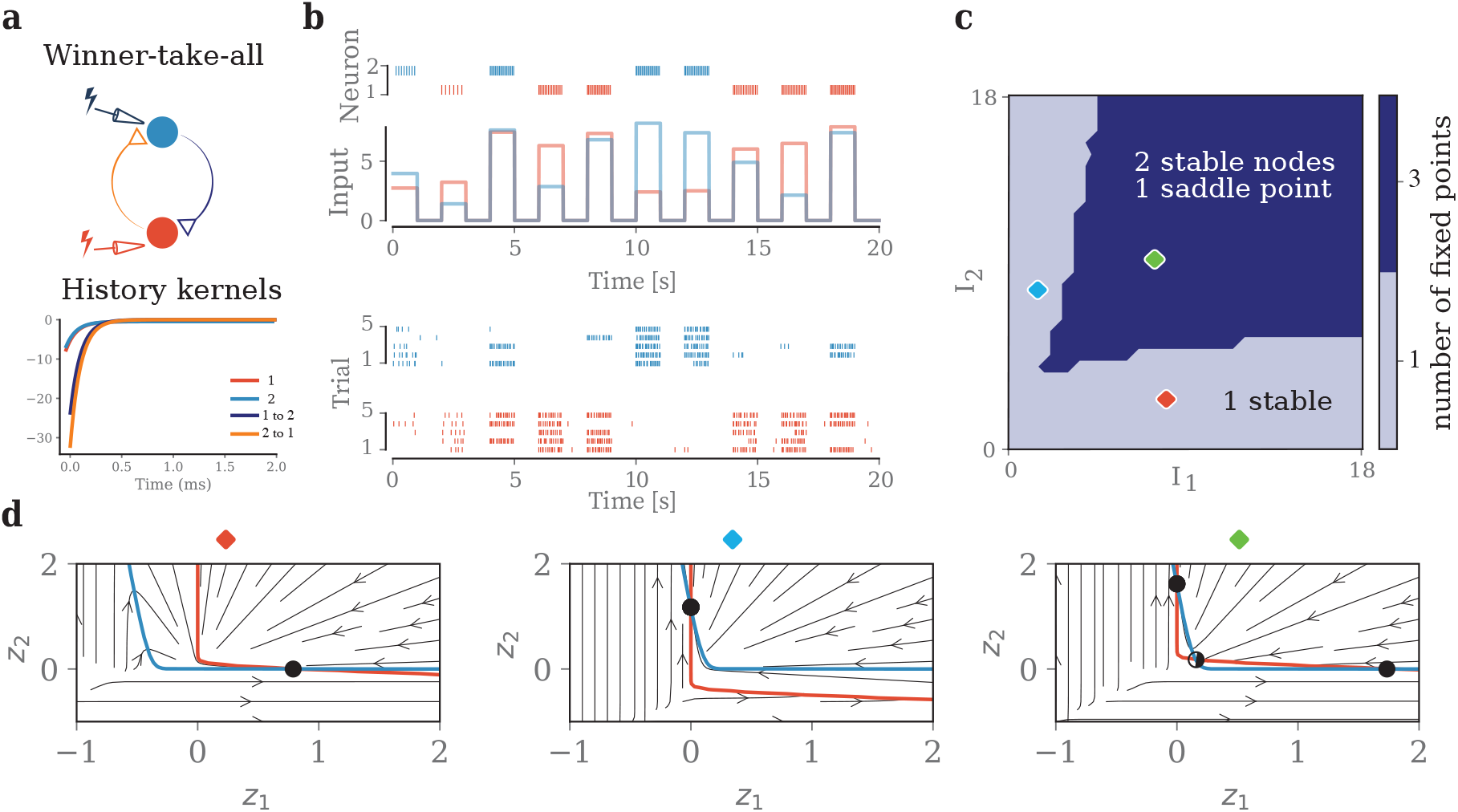
Dynamical repertoire of neuronal-network NH-PDMP models, Winner-take-all: (**a**) Top: Network of two Izhikevich neurons for winner-take-all dynamics. Bottom: history kernels of the estimated NH-PDMP models. (**b**) Top: each Izhikevich neuron receives a distinct input (blue and red lines) resulting in different spiking trains (blue and red vertical lines). Bottom: simulation of five trials of the fitted two-neuron NH-PDMP models, showing winner-take-all dynamics, except for inputs with small differences. (**c**) Parameter space of the deterministic part of the diffusion SDEs as a function of inputs to each neuron. Colored diamonds indicate different parameter choices for the examples shown in (d). (**d**) State space spanned by the two variables *z*_1_ and *z*_2_ in the diffusion SDEs. Left: larger input to the red neuron, showing that the fixed point moves to high (low) values of the red (blue) neuron state variables. Middle: larger input to the blue neuron, showing that the fixed point moves respectively to high (low) values of the blue (red) neuron state variables. Right: equally large inputs to both neurons, showing bistability with two competing stable fixed points.

Wave propagation was simulated using five Izhikevich neurons connected in a feed-forward manner (Figure 6(a, b)). Any sufficiently large input to the first neuron in this chain resulted in the activation of downstream neurons (Figure 6(d) top). We conducted multiple simulations with varying input current magnitudes to each neuron and collected extensive spike time data from the simulated network (Methods). We estimated NH-PDMP models where both the auto- and cross-history kernels were specified by four exponential functions. The true connectivity matrix (Figure 6(b)) was well captured by the fitted parameters (Figure 6(c)), and simulations of the estimated NH-PDMP models with inputs given only to the first neuron triggered activity in downstream neurons as in the Izhikevich network (Figure 6(d) bottom). We then identified the fixed points of the corresponding diffusion SDEs for incrementally increasing inputs to each neuron. We found one 20-dimensional fixed point in the *z* state space for each input to a neuron. The fixed points corresponding to inputs of different amplitudes to the first neuron are expected to have non-zero components along most of the 20 dimensions, reflecting the firing rate increase in most *z* variables when an input is given to the first neuron. Compared to these fixed points, the fixed points corresponding to inputs to the second neuron take zero values for all the variables *z* reflecting the dynamics of neuron 1, as this neuron never fires for an input to neuron 2. Similar logic should apply to subsequent neurons. To show this, the fixed points were projected onto the first principal components (PCs) using principal component analysis (PCA) (Figure 6(e)). As expected, most variance is explained by the first PC for inputs to neuron 1 (red), by the second PC for inputs to neuron 2, etc. We quantified this by computing the position of the fixed points as a function of the PC (Figure 6(f)). For each input magnitude, the fixed point corresponding to the neuron receiving the input indeed had a high component in the corresponding PC. The subspace projections revealed that the organization of the fixed points in the state space reflect the connectivity of neurons in this simple feed-forward network.

**Figure 6.**
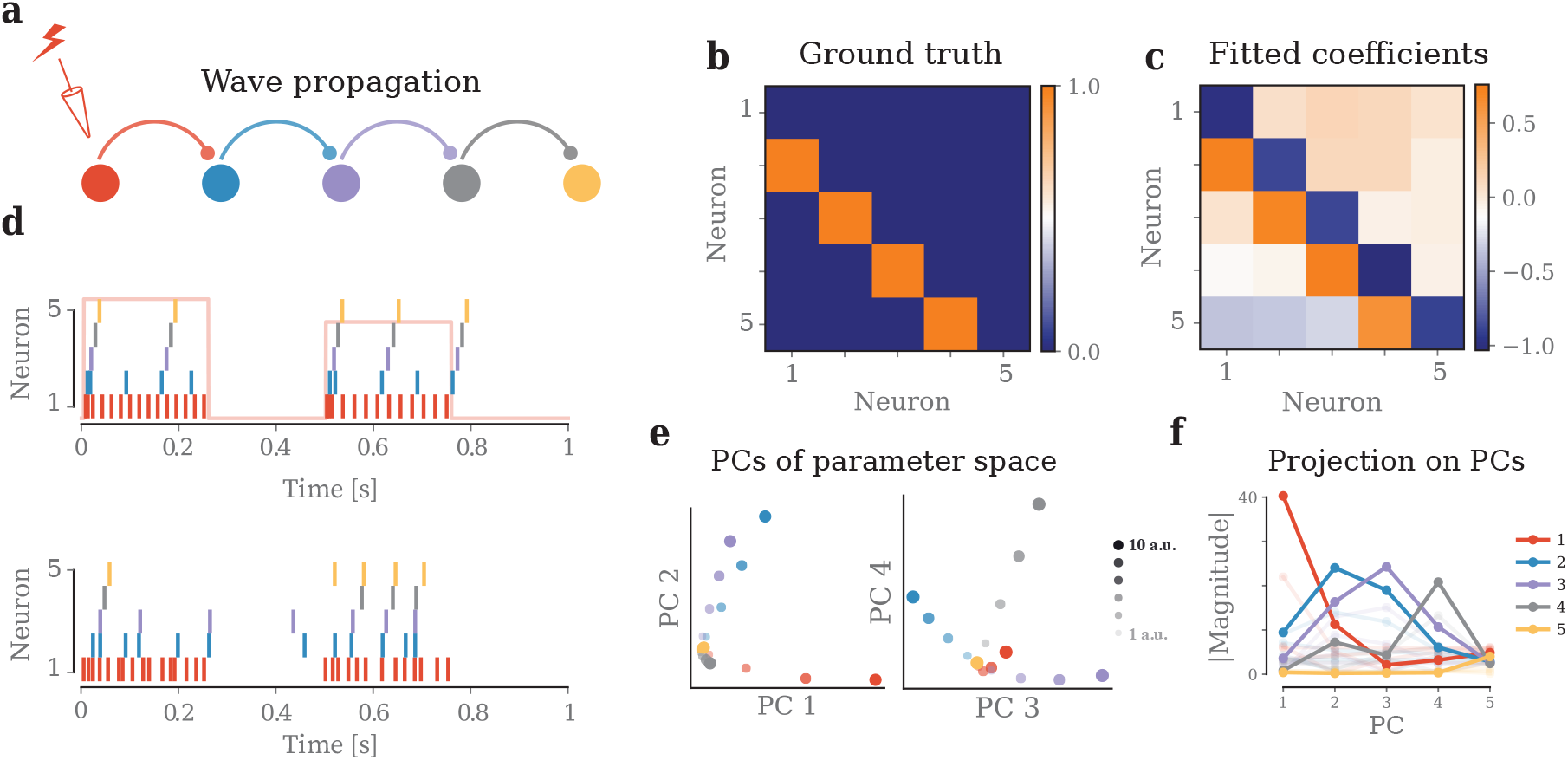
Dynamical repertoire of neuronal-network NH-PDMP models, Wave propagation: (**a**) Network of five Izhikevich neurons coupled in a feedforward configuration (Methods) to support traveling-wave dynamics. (**b**) The feedforward network was specified by the ground-truth connectivity matrix. (**c**) Mean value of estimated parameters across the three cross-history parameters for the fitted NH-PDMP models. (**d**) Top: inputs to the first neuron (red) trigger a spiking train sequence in the downstream neurons. Bottom: simulation of one trial of the fitted five-neuron NH-PDMP model, with a clear spiking traveling wave in the downstream neurons. (**e**) Fixed points of the model projected on the first four principal components of the state space. Different circle sizes indicate the fixed points corresponding to increasing input values to each neuron. (**f**) Magnitude of the fixed-point positions along each PC axis for the highest input value (full line) and other input values (shaded lines). Inputs to neuron earlier in the sequence correspond to high magnitude on lower PCs, indicating their higher impact on the network dynamics.

### Dynamical regimes of data-driven neuronal-network NH-PDMP models fitted to human and nonhuman neuronal recordings

We finally applied our approach to human and nonhuman primate neuronal recordings during two different tasks in order to reveal the neuronal population dynamics underlying specific cognitive and sensorimotor behaviors. As a first application, we analyzed the dynamical properties of the neuronal population activity and their relationship to behavior during a motor control task. We used neuronal recordings from a microelectrode Utah array (MEA) implanted in the primary motor cortex of a nonhuman primate (macaque) during a standard instructed-delay center-out reaching task with eight target directions [31] (Figure 7(a)). Here, we focused on the one-second reaching period following the go cue, and analyzed 115 identified single-units (spike-sorted neurons) over this time period (Figure 7(b) top). To select the neurons that were tuned to the target direction during the task (Figure 7(a)), we first estimated a NH-PDMP model with inputs to each neuron (Methods). Specifically, tuning to direction was captured by the input model *I*_*i*_(*t*) = *β*_*I*,*i*,1_ cos(*θ*_*r*_) + *β*_*I*,*i*,2_ sin(*θ*_*r*_), where *θ*_*r*_ ∈ [−*π, π*] is the direction of the center-out target in the r-th trial (Methods). The models also included the auto-history kernel with a single exponential function, but no cross-history interactions. We selected the twenty neurons with the highest tuning to movement target according to the norm || *β*_*I*_ || (Figure 7(c)). Previous work has shown that population of neurons in the motor cortex encode the task rotational symmetry of the target [32]. Next, we fitted neuronal-network NH-PDMP model using only these 20 neurons (Figure 7(b) bottom). Specifically, this time, the models also included the cross-history interactions, based on one exponential function (Figure 7(d)). We then analyzed the dynamics of the corresponding diffusion SDE (Figure 7(b) bottom). We found that each of the eight targets for the reaching movements corresponded to a single stable fixed point in the parameter space. When projecting the twenty-dimensional state space of the diffusion SDE variables ***z*** onto a two-dimensional space using PCA, we observed that these eight fixed points formed a one-dimensional closed-orbit manifold, in which nearby fixed points corresponds to nearby target directions on the screen (Figure 7(e)). To test whether this organization with rotational symmetry generalized to other direction targets, we additionally used 72 new targets directions sampled uniformly for the [−*π, π*] interval as input directions to the model. The corresponding fixed points formed a similar one-dimensional manifold, and respected the same rotational symmetry (Figure 7(e)). Our analysis thus showed that the rotational symmetry present in the task and encoded by the neuronal population activity is analogously reflected in the dynamical properties of the fitted neuronal-network NH-PDMP models and the corresponding diffusion SDEs.

**Figure 7.**
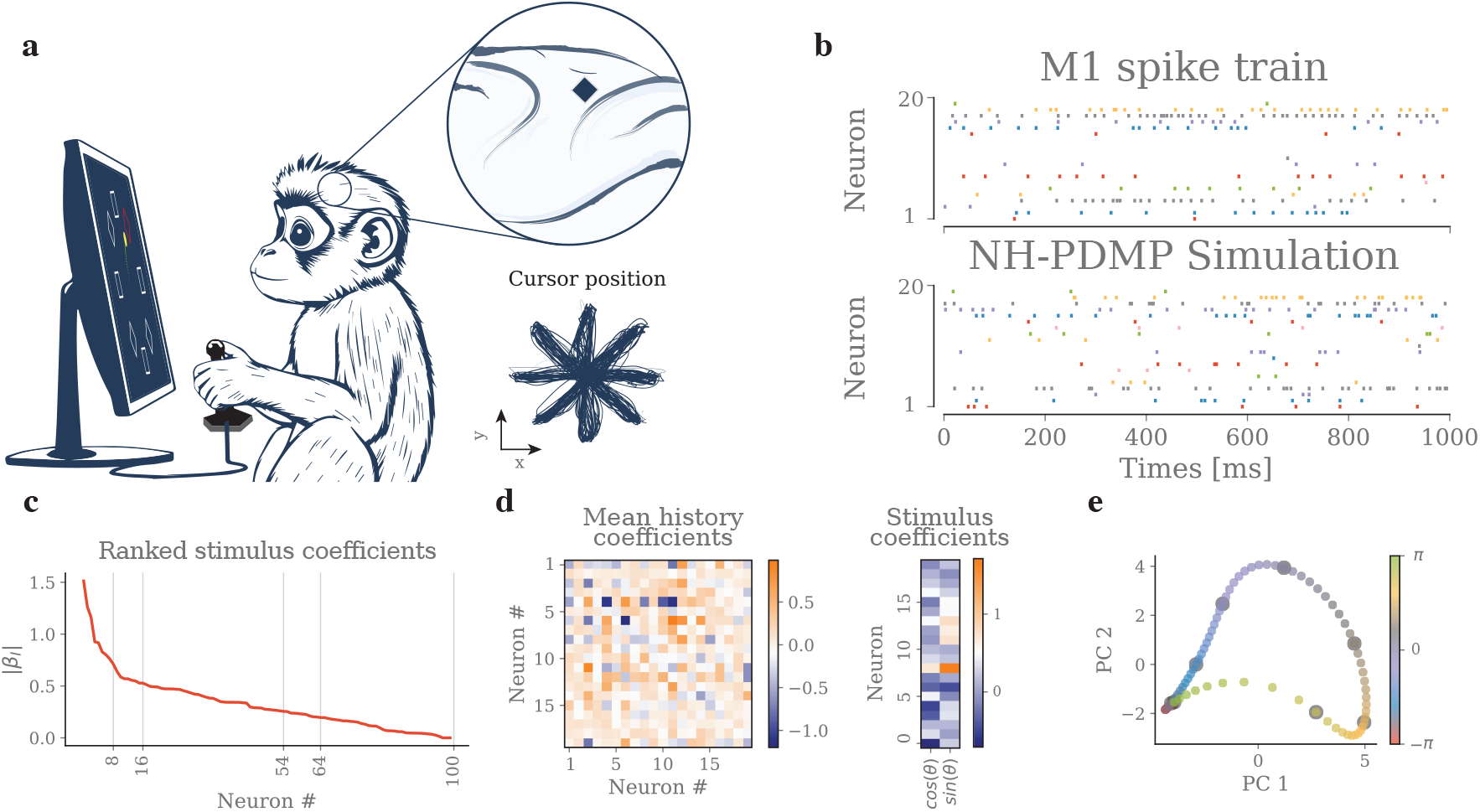
Dynamics of neuronal-network NH-PDMP models fitted to nonhuman primate neuronal population recordings: (**a**) Center-out reaching movement task. Spiking activity from a population of single-units was recorded via a microelectrode array implanted in the primary motor (M1) cortex of a macaque monkey during in a center-out-reach task (top right). Bottom right, cursors position across trials. (**b**) Top: spike train raster of 20 out of 115 sorted M1 single-units during the task. Time zero corresponds to the go-cue onset. Bottom: simulation of the corresponding 20-neuron network NH-PDMP model fitted to the data. The model included an input corresponding to the target direction in each of the corresponding center-out reach trial (as shown in A). (**c**) Recorded neurons were ranked according to their tuning to target direction as captured by the norm of the input parameters || *β*_*I*_ ||. (**d**) Parameters for cross-history history kernels (left) and stimulus contribution (right). (**e**) The eight target directions corresponded to eight fixed point in the 20-dimensional diffusion SDE state space, respectively. Their positions along the first two PCs of the 20-dimensional state space are shown here as black circle. The fixed points corresponding to 72 new simulated target directions are shown as well. The fixed points are organized along a one-dimensional closed-orbit manifold, reflecting the rotational symmetry present in the task.

As a second application, we focused on the neural dynamics underlying neural encoding of phonemes during speech processing [28, 29]. We analyzed recordings from a MEA implanted in the anterior superior temporal gyrus of a 31-year-old participant with pharmacologically resistant epilepsy (Figure 8(a)). The participant performed an auditory semantic categorization task, during which he indicated by button press whether the spoken words represented objects smaller or larger than a foot in any dimension. The experiment involved 410 nouns, encompassing a range of objects, animals and body parts of varying sizes. 176 single-units were spike-sorted from the MEA recordings. There we sought to identify neurons responding only to the phonological content of the stimuli. An NH-PMDP model was estimated for each unit individually, excluding cross-history interaction terms and used the ground-truth phonemes’ identities as exogenous input (Figure 8(b)). We selected the 20 neurons with the highest tuning (parameters) to phoneme onsets and estimated an NH-PDMP model on selected neurons, this time incorporating also the cross-history interactions and using 36 exogenous inputs (corresponding to phoneme identities for both vowels and consonants) (Figure 8(c)). Using the corresponding diffusion SDEs, we identified the fixed points of the resulting 20-neuron network that corresponded to each vowel input. A projection of the state space of the diffusion SDE variables ***z*** into a lower-dimensional space using PCA revealed a spatial organization. Specifically the fixed points that related to vowels with a first formant frequency below 500 Hz were spatially segregated from those with a first formant frequency above 500 Hz (Figure 8(d) left), confirming previous results [29]. No similar separation was observed for vowels based on their second formant frequency above or below 1.5 kHz (Figure 8(d) right) [29].

**Figure 8.**
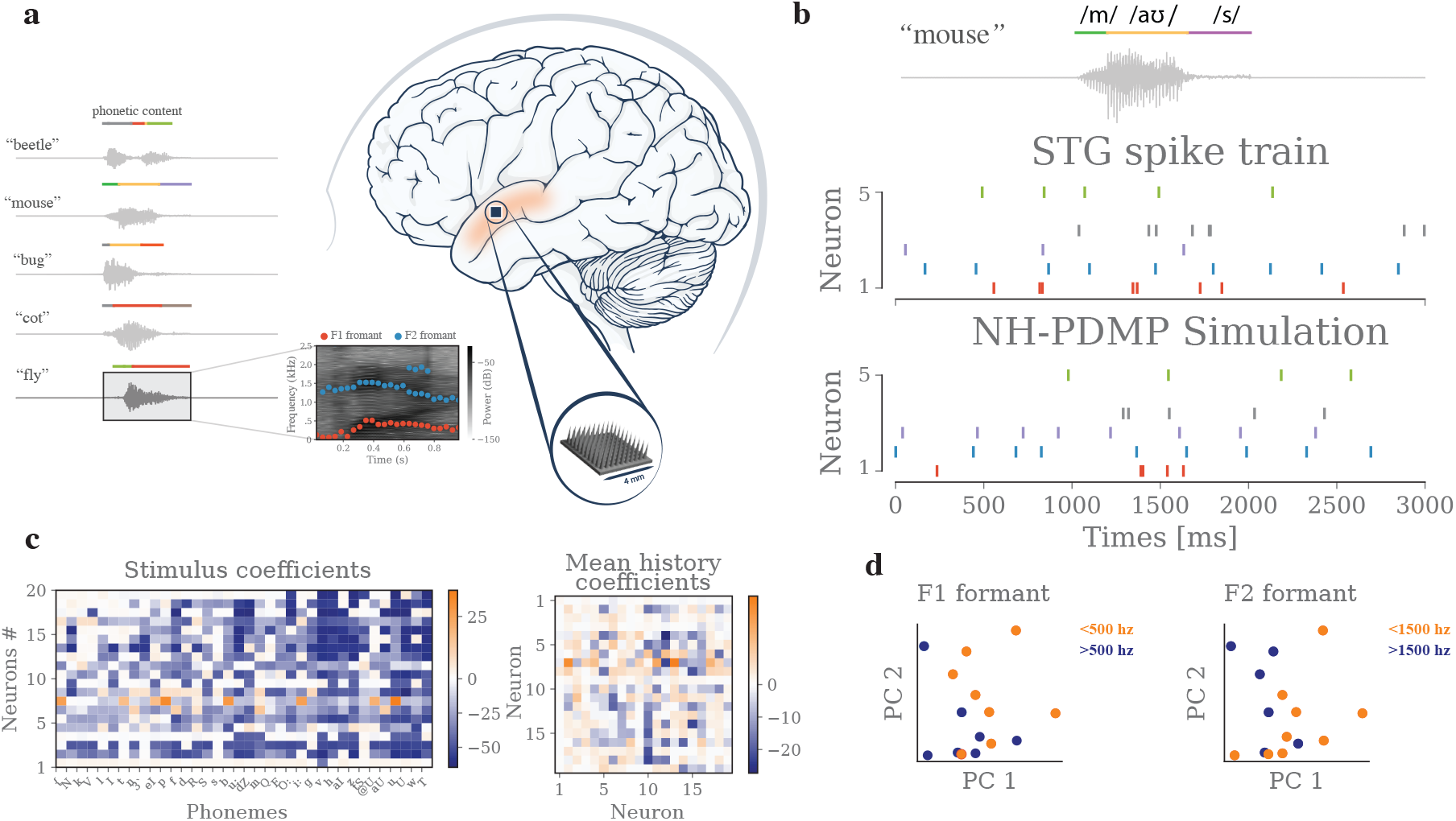
Dynamics of neuronal-network NH-PDMP models fitted to human neuronal population recordings: (**a**) Speech processing task. Left: speech waveforms, and phonemic content for five example words used in the semantic categorization task [28, 29]. Right: Microelectrode array recordings located in the anterior superior temporal gyrus (aSTG). Brain picture in panel A adapted from NIH BIOART Source [30]. (**b**) Estimation of NH-PDMP models. The input to the NH-PDMP models consisted of the phonemes present in the heard word, here “mouse” (top). Spike train recordings of five neurons for one trial for the word “mouse” (middle). Corresponding simulation of the estimated NH-PDMP models (bottom). (**c**) Parameters for the cross-history kernels (left) and inputs (right). (**d**) The 15 fixed points in the 10-dimensional diffusion SDE state space that corresponded to each vowel input were projected into the first two PCs of the state space. Left: a separation of the fixed points is observed on the basis of the first formant of vowels (frequency below or above 500 Hz). Right: no such separation is observed for the second formant of vowels (frequency below and above 1.5 kHz).

## Discussion

We present a new approach to reveal the nature and repertoire of single-neuron and neuronal-network spiking dynamics based on two main steps. First, we equivalently represented nonlinear Hawkes processes as nonlinear-Hawkes piecewise deterministic Markov processes (NH-PDMP) to model neural dynamics. These NH-PDMP models consist of system of differential equations with stochastic jumps. Second, we applied a diffusion approximation to the jump processes to obtain a continuous stochastic differential equation (diffusion SDEs) approximation of the original NH-PMDP models. This approximation enabled us to apply methods from nonlinear dynamical system analysis, thereby providing quantitative and geometric insights into both the stability and dynamics of the spiking activity of populations of neurons. For a single neuron, we derived a closed-form solution for the fixed point of the deterministic part of the diffusion SDEs, and analytically predicted their existence and stability. The full stochastic diffusion SDEs for a single neuron were solved using a moment-closure approach, whose solutions accurately matched the exact continuous-time simulations of the NH-PDMP models performed with an algorithm specifically designed for it. These analyses were used to uncover the dynamical properties that underlie the canonical repertoire of single neurons including type I and type II excitation, tonic and phasic spiking and bursting. Our approach successfully extended to network of neurons, and when applied to winner-take-all and wave propagation examples, it revealed the dynamical properties underlying the different regimes. We finally applied these analyses to neuronal population recordings in human and nonhuman primates during speech processing and center-out reach movement tasks, respectively, and showed that the dynamical properties of the diffusion SDEs of the estimated NH-PDMP models recovers the characteristic of the stimuli that were encoded by the population of neurons.

Given that the neuronal NH-PDMP models examined here are far from satisfying the strict conditions typically required for these approximations [21], the adopted diffusion approximation works remarkably well. As stated earlier, we conjecture that this efficacy results from smooth history kernels and the not-too-sparse spiking activity. We also note our analyses focused on the short-term dynamics in the single-neuron and neuronal-network dynamics, rather than on long-term time-invariant distributions. Accordingly, we also defined stability in two broader ways, in contrast to stricter definitions based on stability in variation [7, 9], for example.

The diffusion approximation operates over time and preserves the full dimensionality of the population dynamics. This stands in contrast to mean-field approaches, which use a diffusion approximation across neurons to derive the dynamics of the mean of the neuronal population activity [33, 34]. Instead, our approach enables the identification of complex encoding schemes that may depend on the dynamics of the covariance and higher moments of the population of neurons. For example, it can be combined with analytical or dimensionality reduction methods to uncover the mechanisms by which complex nonlinear dynamics encode for behavior in low-dimensional subspaces [29, 35–37]. Another investigation avenue offered by our approach is the analysis of small and well-identified microcircuits of neurons, such as the central pattern generator circuits in lobsters [38] or retinal circuits [39]. Nevertheless, the dynamics we analyze reflect the functional network dynamics contributing to a specific behavior or cognition. Linking functional network dynamics with the underlying structural microcircuitry necessitates additional analysis [40].

The diffusion approximation offers a novel perspective on the dynamics of single-neuron and neuronal-network processes and models. Instead of directly analyzing the neuronal voltage supporting subthreshold and spike dynamics, it focuses on the temporally smoothed presynaptic inputs and the directly related conditional intensity function (i.e. the instantaneous spiking rate). The dynamical repertoire underlying distinct spiking behaviors operates in the “rate” space of the diffusion SDE, which differs from the voltage space. For instance, the tonic spiking regime corresponded to the displacement of a single stable fixed point to a higher firing rate, rather than a saddle-node bifurcation in the voltage space. This approach enabled to identify the way in which the estimated NH-PDMP models captured diverse dynamical behaviors, as demonstrated through simulations and human and nonhuman primate recordings. It also permits the analytical prediction of the region of instability of single and network of neurons. Additional strategies can be used to adjust the weights of the history kernels of divergent models and stabilize their dynamics with minimal alterations [10, 12].

The rate perspective offered by the diffusion SDEs is analogous to that of commonly used recurrent neural networks (RNNs) based on mean firing rate models, which directly estimate the activity rate for individual neurons [35, 41]. However, a key distinction between these two models lies in their respective approach to spike representation during the fitting process. These RNN models require to fit an arbitrary smoothing function to the neuronal spike trains. In contrast, estimation of NH-PDMP models is directly based on the likelihood of the actual spike train data, allowing for high temporal precision depending on the selected time scales of the basis functions. The diffusion approximation then introduces a degree of smoothing, the accuracy of which is not arbitrary but depends on the firing rate of the neuron. Additionally, under the choices made here for the type of nonlinearity and the parametric form of the NH-PDMP models, parameter estimation is a convex optimization problem that guarantees a unique solution if the solution exists, in contrast to the fitting of commonly used RNN rate models. (Solution existence issues can be easily dealt with regularization.) A current limitation of the use of PPGLM framework to fit NH-PDMP models is that the time constants of the chosen exponential function bases need to be set beforehand via some exploration. (They are not loglinear in the conditional intensity functions and cannot thus be estimated in the above PPGLM framework; see Methods for details). We hope to address this issue in future studies. Nevertheless, our analytical approach to very more general theoretical or data-driven NH-PDMP models is not by itself limited by this constraint.

With the continuing increase in the number of recorded neurons, it becomes critical to develop new methods to synthesize the organizational principles of their dynamics. NH-PDMP models together with the diffusion approximation approach proposed here offer an effective alternative for studying the dynamics of networks of neurons. Our approach allows extracting tractable equations while preserving the high-dimensional nature of the neural data, opening the way for applying diverse dynamical analysis and dimensionality reduction methods [2, 37, 42, 43].

## Methods

### Stability and dynamics of single-neuron NH-PDMP models

#### Fixed points, stability and bifurcations of the deterministic skeleton of the diffusion SDE

The fixed points in the deterministic part of the diffusion SDEs (Eq. 4) for single nonlinear Hawkes processes with *L* basis functions was analytically derived by setting the temporal derivatives to 0, i.e. ***z***^*∗*^ = ***T*** 𝟙*λ*^*∗*^ with *λ*^*∗*^ = exp *β*_0_ + ***β***^*T*^***z***^*∗*^ + *I* the conditional intensity function at the fixed points ***z***^*∗*^ for a given input *I*. Moving terms and multiplying by −***β***^*T*^ gives

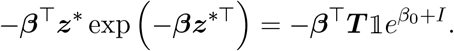

The closed-form solution of this equation is based on the Lambert function 𝒲_*{*0,1*}*_

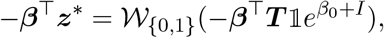

and the fixed point are given by

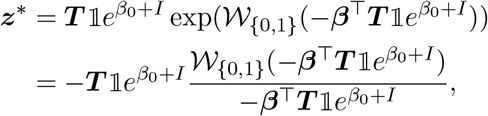

resulting in (Eq. 6). As described in the Results section, there are only zero, one, or two fixed points depending on the chosen parameters ***β, T***, *β*_0_, and *I* (Figure 2(A)).

The linear stability of each fixed point is obtained by computing numerically the eigenvalues of the Jacobian matrix **J** = −***T*** ^*−*1^ + 𝟙***β***^*T*^*λ*^*∗*^. For one and two exponential kernels, the stability of the system is also determined analytically from the trace and the determinant of the Jacobian matrix [44]. The nature of the bifurcations is determined numerically using BrainPy continuation and bifurcation software [45].

#### Moments of the diffusion SDEs

To analyze the full diffusion SDEs, including the stochastic contributions, we performed a moment expansion and closure using the moment equation (from Itô’s lemma) to compute the evolution of the system’s first moments ⟨***z***⟩ and second central moments (covariance matrix) **Σ**= ⟨(***z*** −⟨***z***⟩)(***z*** −⟨***z***⟩)^*T*^⟩ [23] (for notational convenience, we omit the time dependency for ⟨***z***⟩(*t*), **Σ**(*t*), and ⟨*λ*⟩(t)):

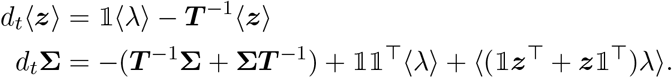

The two nonlinear expressions ⟨*λ*⟩ and ⟨ (𝟙***z***^*T*^ + ***z***𝟙*^T^*)*λ*⟩ can be evaluated by deriving analytically these two expectations using Gaussian moment closure.

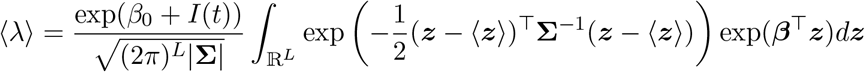

By change of variable 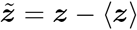we obtain

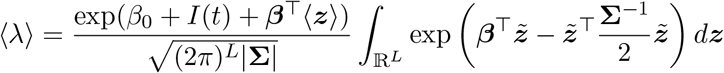

Using a known identity for multivariate integrals, and simplifying, then gives

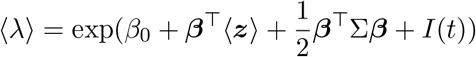

Similarly we can compute

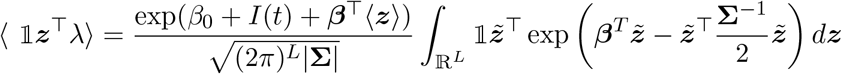

Using known identities for multivariate Gaussian integral, the expression for ⟨*λ*⟩, and simplifying, then gives

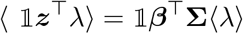

and the final result

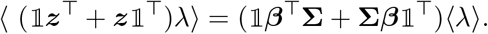

### Stability and dynamics of neuronal-network NH-PDMP models

#### Fixed points, stability, and bifurcations of the deterministic part of the diffusion SDE

Setting the temporal derivatives of the deterministic part of the diffusion SDEs (Eq. eq. (5)) to zero leads to the fixed point equation for the network of NH-PDMP models, ***z***^*∗*^ = ***T*** 𝟙***λ***^*∗*^ where 𝟙 is now a block diagonal matrix of size *JL* × *J* where each block is 𝟙 = (1, …, 1)^*T*^ with *L* ones, and ***λ*** = (*λ*_1_, …, *λ*_*J*_)^*T*^ is a vector of size *J*. No closed-form solution exists for this equation which we solved numerically. The corresponding Jacobians and bifurcations are established numerically using the BrainPy library [45].

#### Moments of the diffusion SDE

Similarly to the one neuron case, moment expansion and closure can be performed analytically to obtain the first moments ⟨***z***⟩ and second central moments (covariance matrix) **Σ**= ⟨(***z*** − ⟨***z***⟩)(***z*** − ⟨***z***⟩)^*T*^⟩, which are now of size *JL* and *JL* × *JL*, respectively, are [23]:

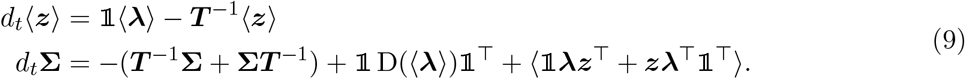

Deriving the Gaussian expectations as in the single neuron case and accounting for the fact that ***B*** is now a matrix and ***λ*** a vector, we find:

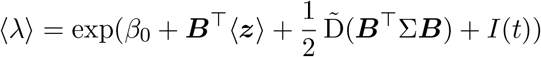

and

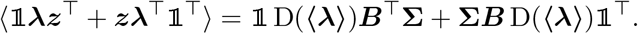

can also be evaluated analytically from the Gaussian expectations.

### Exact continuous-time simulation of single-neuron and neuronal-network NH-PDMP models

Most previous approaches for exact continuous-time simulation of NH-PDMP models have relied on thinning algorithms [16]. Here, we develop an exact simulation algorithm based in part on the time-rescaling theorem. The algorithm for simulation of single-neuron and neuronal-network NH-PDMP models has two main components.

First, the algorithm applies the time-rescaling theorem [46] to the simulation of point processes as previously suggested [19, 20]. Briefly, any stochastic realization of a single-neuron NH-PDMP model consists of a sequence of spike times {*t*_1_, *t*_2_, *t*_3_, … } under a given conditional intensity function (CIF) model *λ*(*t*|H_*t*_) (eq. (2)). By the time-rescaling theorem, the rescaled inter-spike interval (ISI) times

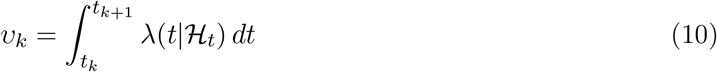

are independent and exponentially distributed with mean 1. In other words, the above time rescaled process corresponds to a homogeneous unit rate Poisson process. Thus, to simulate the original NH-PDMP model, one first samples a long enough sequence of unit mean exponential ISIs {*υ*_*k*_}. Next, for any given *t*_*k*_, one solves eq. (10) for next spike time *t*_*k*+1_ of the NH-PDMP model being simulated. The integral is solved via numerical methods with care regarding the choice of step size, especially when dealing with unstable NH-PDMP models. We start at time *t* = 0 and explore different initial conditions for the ***z*** state variable in the NH-PDMP model.

Second, the algorithm takes advantage of the deterministic linear dynamics in between the occurrence of stochastic jump (spike) events in the PDMP representation. Briefly, this deterministic linear dynamics allows us to easily evolve the state ***z*** and the CIF in the integrand when solving eq. (10) via a matrix exponential operator. Specifically, in between any two stochastic jumps, the deterministic part of the SDEs eq. (2) can be solved via

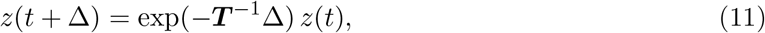

where ***T*** = diag(***T***_1_, …, ***T***_*J*_) is the appropriately defined state matrix according to eq. (2) and Δ here denotes the time step when numerically solving the integral equation eq. (10). (Given the initial conditions, one can start at *z*(0).) Thus, the above allows the time evolution or update of *λ*(*t*|ℋ _*t*_) using eq. (2) at any time point when solving the integral equation (eq. (10)).

The exact continuous-time simulation algorithm for single-neuron based on the time-rescaling theorem can be easily extended to multidimensional systems [20], including in particular the neuronal-network NH-PDMP models examined in this study. In this case, the conditional intensity functions *λ*_*i*_(*t*|ℋ_*t*_) depend also on the the spiking history of other neurons in the population and are updated according to the neuronal-network SDEs with stochastic jumps (eq. (2)). As before, a long enough sequence of ISIs are sampled from a unit mean exponential distribution and the integral equation to be solved now corresponds to

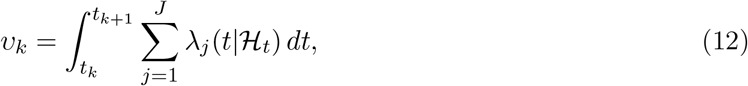

where the sum in the integrand corresponds now to the population intensity. In other words, one is sampling a sequence of spike times that occur in the population. Because in continuous-time simultaneous events in two or more neuron within this population point process have measure zero [47], each new spike time needs to be probabilistically assigned to a unique neuron in the population.

This assignment is done by sampling from a multinomial distibution where each spike event can be assigned to a specific neuron *m* with probability

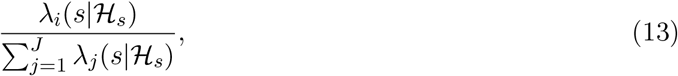

evaluated at the solution *s* = *t*_*k*+1_ of eq. (12). As for the single-neuron model case, the deterministic time evolution of the SDEs eq. (2) in between stochastic spikes can be implemented via the matrix exponential operator eq. (11), with the state matrix ***T*** ^*−*1^ now corresponding to the appropriate matrix of the full SDEs eq. (2).

### Simulation of single-neuron and neuronal networks based on Izhikevich canonical models

To examine how our approach captures the dynamical repertoire of complex neuronal spiking patterns, we used single-neuron and neuronal networks based on the Izhikevich quadratic models [6, 25]. The Izhikevich model is a two-dimensional model, where the variable *v*_*i*_(*t*) tracks the membrane potential of neuron *i*, and the variable *u*_*i*_(*t*) acts as a recovery variable

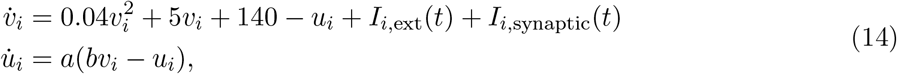

where *I*_*i*,ext_(*t*) denotes the external stimulus of neuron *i* and *I*_*i*,synaptic_ represent the total synaptic input from all other neurons to neuron *i*. A hard membrane potential threshold determines the spike resetting conditions:

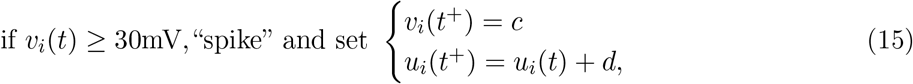

where *t*^+^ is the time step following the spike occurring at time *t*, with the membrane potential reaching the threshold of 30 mV. The dynamics of the neuron is characterized by the dimensionless parameters (*a, b, c, d*).

The network interactions are introduced via the total synaptic input *I*_*i*,synaptic_ as the sum of the synaptic currents *s*_*j*_ from all other neurons in the network, each weighted by the corresponding synaptic weight

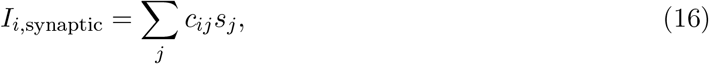

where *c*_*ij*_ is the weight of the synapse from neuron *j* to neuron *i*. The synaptic currents *s*_*j*_ are modeled as an exponentially decaying function of time 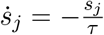 with *τ* the decay time constant, and are increased by a constant *s*_*j*_(*t*^+^) = *s*_*j*_ + *g*_*syn*_ each time the corresponding neuron *j* emits a spike.

### Single-neuron canonical models

We used multiple types of Izhikevich canonical models and adopted previously reported parameters (table 1). These parameters were chosen based on a previous study [5]. To generate the training set used to fit the NH-PDMP model, for each canonical behavior, we used 10 successive 0.6 second long step inputs separated by equal intervals of 0.4 seconds. For type I and type II neurons, the amplitude of the step current gradually increased, while for all other neurons it was kept constant (amplitudes given by *I* column in 1). To generate the test set, we used 20 trials of one second simulation time.

We used one (tonic and phasic spiking and bursting) or three (class I and II) exponential kernels for fitting and one second of simulation time for both NH-PDMP and Izhikevich models as the test set. All simulations of Izhikevich models in this and following sections used a forward-Euler scheme with a time step size of 0.1 ms.

### Winner-take-all network

To simulate the winner-take-all behavior, we used two coupled Izhikevich neurons with inhibitory coupling by defining the connectivity matrix as 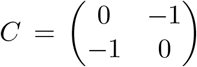 . To generate the training set for estimating the NH-PDMP model, we excited both neurons with one second long step inputs separated by intervals of one second and with different amplitudes, which were sampled from a uniform distribution between 2.5 pA and 7.5 pA. The neuron with the highest input dominates the competition and inhibits the other neuron. Simulations were 20 second long. For the test set, we used ten step inputs and 20 seconds of simulation for both Izhikevich neuron and NH-PDMP simulations.

### Propagating wave network

To capture the behavior of the propagation wave, we used five Izhikevich neurons coupled in a feedforward fashion by connectivity *c*_*i*,*j*_ = *δ*_*i*,*j*+1_, where *δ*_*i*,*j*+1_ is the Kronecker delta function, which is equal to 1 if *i* = *j* + 1 and 0 otherwise. 200 step inputs were used, with one second long step inputs separated by intervals of one second and with amplitude sampled from the uniform distribution between 2 pA to 20 pA. Each input was fed to only one neuron, which was selected using linearly spaced probabilities, starting from 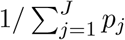 for the first neuron in the sequence and decreasing to 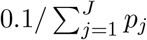 for the last neuron in the sequence. Simulations were 20 seconds long. For the test set, we used two step inputs and one second of simulation for both Izhikevich neuron and NH-PDMP simulations.

### Human and nonhuman primate neuronal recordings

#### Center-reach-out dataset

We used an open-access dataset recorded from the primary motor (M1) and dorsal premotor (PMd) areas of the motor cortex in a nonhuman primate (macaque monkey) [31]. Specifically, we employed data recorded from one monkey (Monkey M), trained to execute a standard center-out reaching task with eight peripheral targets distributed uniformly on a circle with an 8 cm radius on the screen. Each trial began by the monkey moving a pointer to the center of the screen using a joystick, maintaining this position for 0.5 to 1.5 seconds. Following this holding period, one of the eight targets appeared on the screen, initiating a delay period of 0.5 to 1.5 seconds. The monkey then received an auditory go cue, prompting the center target to disappear. The monkey was required to move the joystick to reach the presented target within 1 second and maintain this position for 0.5 second.

During the experiment, neural activity was recorded using an implanted 96 microelectrode Utah array. Neuronal spike times were detected using spike sorting methods, identifying 66 single units from the PMd and 34 single units from M1.

For this study, we estimated NH-PDMP models from the spike trains from the go cue time up to one second of spike data. To reduce the number of neurons for fitting and analysis, we initially fitted all neurons, taking into account only an exogenous input (the direction of the center-out target) and auto-history effects but not cross-history interactions. The tuning to direction was modeled as *I*_*i*_(*t*) = *β*_*I*,*i*,1_ cos(*θ*_*r*_) + *β*_*I*,*i*,2_ sin(*θ*_*r*_), where *θ*_*r*_ ∈ [−*π, π*] is the direction of the center-out target in the rth trial (see below for details on the statistical estimation). The auto-history kernels consisted of a single exponential with a time constant of 0.01 s. Neurons were then ranked based on their tuning to direction (input parameters), and the top 20 neurons with the highest tuning were selected. Next, we estimated neuronal-network NH-PDMP models to these 20 neurons. Cross-history kernels also consisted of a single exponential with the same time constant as for the auto-history kernels. Next, we used the diffusion approximation with the obtained parameters from the NH-PDMP fitting. We then used the BrainPy library to identify the high-dimensional fixed point of this 20-neuron network across various *θ* values ranging from [−*π, π*]. To see if there’s an organizational structure between these obtained fixed points, we used PCA to reduce the dimensionality of the state space.

### Speech task dataset

A 31-year-old male participant with drug-resistant epilepsy underwent semi-chronic electrode implantation at Massachusetts General Hospital as part of a surgical evaluation [28, 29]. His seizures, originating from the mesial temporal region, led to the resection of the left anterior temporal lobe, parahippocampal gyrus, hippocampus, and amygdala. Post-surgery, he became seizure-free without significant language deficits. The participant provided informed consent for the study, which was conducted under ethical guidelines. A microelectrode array (Blackrock Microsystems) was implanted in the left anterior superior temporal gyrus (aSTG) to record single-unit action potentials. The 4 × 4 mm array contained 100 electrodes (96 active), each 1.5 mm long with 20-µm platinum tips spaced 400 µm apart. Neural data were recorded at a 30 kHz sampling rate using a Blackrock NeuroPort system. The array was positioned in the superior temporal gyrus based on clinical requirements and was removed during surgery. Surrounding tissue was excised, stained, and examined, confirming electrode tips near the border of cortical layers III and IV, with normal adjacent tissue.

The participant performed several auditory tasks to evaluate speech and nonspeech sound processing. In the auditory semantic categorization task, the participant was instructed to press a button upon hearing spoken words denoting objects or animals larger than one foot in any dimension. The words, spoken by a male, were standardized to 500 ms in duration, with a 2200 ms interval between each. There were 800 trials, consisting of 400 unique words and 10 repeated words that appeared 40 times each. To prevent bias in the model, we excluded all repeated trials from the analysis. Spikes were extracted using an unsupervised spike detection algorithm [48].

To identify neurons most responsive to speech content, we initially estimated single-neuron NH-PDMP models (i.e. without incorporating interactions across all 176 neurons), using the spiking activity and all 36 phonemes from 400 words as external inputs (see below for details on the statistical estimation). Neurons were ranked based on the absolute sum of their stimulus parameters, and the 20 neurons with the highest parameters were selected. A subsequent neuronal-network NH-PDMP model was then fitted, allowing it to capture cross-coupling interactions. Using the fitted history and stimulus parameters in the corresponding diffusion SDE, we computed the fixed point in the 20-dimensional state space corresponding to each vowel given as an input. PCA analysis on the state space including all fixed point showed that fixed points associated with vowels having a first formant below 500 Hz clustered closer together compared to those with a first formant above 500 Hz [28, 29, 49–51].

### Statistical estimation of single-neuron and neuronal-network NH-PDMP models

To estimate NH-PDMP models to the spike events from model simulations or actual neuronal population recordings, we applied the (discrete-time) point-process generalized-linear-model statistical estimation framework [1, 10]. The solution exists and estimated parameters are guaranteed to be unique under appropriate regularization [52–54].

In this context, two main preliminary aspects are considered. First, the choice of nonlinearity. When using this statistical framework for statistical estimation, the time-discretized likelihood function belongs to the exponential family, in particular the Poisson one. In this case, the natural or canonical parameter is the logarithm of the conditional intensity function (CIF). Therefore, modeling log *λ* = *f* (*x*), leads to the exponential nonlinearity choice: *λ* = exp[*f* (*x*)]. Additional experimental motivation also exists [55].

Second, NH-PDMP model estimation in the above statistical framework setup requires the free model parameters to be linear in the log *λ* scale. (The actual inputs can enter the model via any nonlinear function, but the free parameters need to be loglinear in the CIF model.) In this way, only certain parametric forms for the history kernels can be used in NH-PDMP models. Specifically, the time constants in the exponential functions are not free parameters to be estimated within the point-process generalized-linear-model framework. They should instead be carefully chosen and set to fixed values before hand to account for the main time scales of interest, similarly to what has been done before with raised cosine and other basis functions [3, 56]. For more details on the above two aspects see [1, 2, 57].

Here, the auto- and cross-history kernels *h*_*ij*_(*t*) in eq. (1) were written as a sum of weighted exponential basis functions

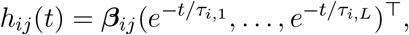

where the *τ*_*i*,*k*_ were fixed (table 2).

The parameters ***β***_*ij*_ and offsets *β*_*ij*,0_ were inferred by maximum likelihood estimation. We used a discrete-time representation of the original neuronal point process by partitioning any of the recording period of length *T* into regular *M* = *T/*Δ time bins, each with size Δ. Here, we set Δ = 1 ms or smaller depending on the dataset, which resulted into binary spike train sequences *y*_*j*,*m*_, *m* = 1, …, *M* . The corresponding data log-likelihood function under a given CIF model with a parameter set Θ is then proportional to

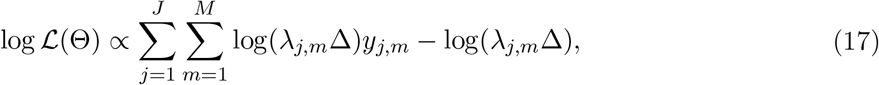

where *λ*_*j*,*m*_ = *λ*_*j*_(*m*Δ | ℋ_*m*Δ_, *I*_*j*_(*m*Δ)). The estimation was performed with the statsmodels Python library [58], while specifying a log link function and the Poisson distribution for the corresponding generalized linear models. To fit the point-process generalized-linear-model parameters on the speech dataset, we used ridge regularization. No regularization was used for models fitted to the center-reach-out dataset.

As a goodness-of-fit test, we applied the time rescaling theorem [1, 19, 59] and used the Kolmogorov-Smirnov (KS) test. The KS test compares the empirical cumulative distribution function (CDF) of the time rescaled sampled inter-spike intervals under the estimated CIF model with the CDF of the exponential distribution with mean 1.

## Competing Interests

The authors declare no competing interests.

## Acknowledgements

We thank Michael Rule for fruitful discussions. This work was funded by Swiss National Science Foundation career grant 193542 (T.P.), Ernst et Lucie Schmidheiny Fondation scholarship (N.M.), National Institutes of Health (NIH), National Institute of Neurological Disorders and Stroke (NINDS), grant R01NS079533 (W.T.), and the Pablo J. Salame Goldman Sachs endowed Associate Professorship of Computational Neuroscience at Brown University (W.T.).

## Notes

### Competing Interest Statement

The authors have declared no competing interest.

